# An ANI gap within bacterial species that advances the definitions of intra-species units

**DOI:** 10.1101/2022.06.27.497766

**Authors:** Luis M. Rodriguez-R, Roth E. Conrad, Tomeu Viver, Dorian J. Feistel, Blake G. Lindner, Fanus Venter, Luis Orellana, Rudolf Amann, Ramon Rossello-Mora, Konstantinos T. Konstantinidis

**Author notes:** Correspondence should be addressed to K.T.K. equal first authors (Luis M. Rodriguez-R, Roth E. Conrad).

## Abstract

Large-scale surveys of prokaryotic communities (metagenomes) as well as isolate genomes have revealed that their diversity is predominantly organized in sequence-discrete units that may be equated to species. Specifically, genomes of the same species commonly show genome-aggregate average nucleotide identity (ANI) >95% among themselves and ANI <90% to members of other species, while genomes showing ANI 90-95% are comparatively rare. However, it remains unclear if such “discontinuities” or gaps in ANI values can be observed within species and thus, used to advance and standardize intra-species units such as strains and sequence types. By analyzing 18,123 complete isolate genomes from 330 bacterial species with at least ten genome representatives each as well as available long-read metagenomes, we show that such a discontinuity exists between 99.2-99.8% (mean 99.5%) ANI. The 99.5% ANI threshold is largely consistent with how sequence types have been defined in previous epidemiological studies but provides clusters with ∼20% higher accuracy in terms of evolutionary and gene-content relatedness of the grouped genomes, while strains should be consequently defined at higher ANI values (>99.99% proposed). Collectively, our results should facilitate future micro-diversity studies across clinical or environmental settings because they provide a more natural definition of intra-species units of diversity.

## Introduction

Discrete, or somewhat discrete (1), ecological, functional or evolutionary units within bacterial species have been recognized for some time. These units have been designated with various terms such as subspecies, ecotypes, clonal complexes, serotypes and strains, among several others [reviewed in (2)]. However, the application of these units has commonly been inconsistent between different taxa and studies, e.g., different marker genes and standards for each marker are used, creating challenges in communication about diversity. If diversity within species is indeed organized in discrete units and these units show similar intra-unit relatedness levels, that is the units are consistent across taxa, this information could be used to standardize unit definition and recognition. These challenges are represented well by the use of clonal complex (CC) and sequence type (ST), two terms that are commonly employed to catalogue intra-species diversity. These terms have been successfully used, especially in medical microbiology and epidemiological studies, to identify an outbreak caused by a specific pathogenic organism (or pure isolate) or groups (complexes) of highly related organisms. A ST is typically defined as a collection of genomes with no nucleotide sequence diversity (zero single-nucleotide polymorphisms) in 6-7 selected genetic loci (3). These loci are typically distributed across the genome to avoid co-selection evolutionary events, and represent fragments (PCR amplicons) of genes shared by most members of a species; that is, core genes. While this definition is pragmatic and operational, it has its own inherent limitations. Most notably, different gene markers are often used for different taxa, and core genes tend to be more conserved than the genome average. Thus, it remains somewhat speculative how similar (or not) organisms of the same ST may be in the rest of their genome, and this may also depend, at least partly, on the exact loci used in the analysis since different genes often evolve under varied selective constrains. Several recent efforts have also employed all core genes in the genome, providing a different (higher) resolution level compared to earlier efforts with 6-7 genes, which, nonetheless, makes isolate-typing results based on different sets of genes not directly comparable (4, 5). On the other hand, CCs are defined as closely related STs (complexes) based on phylogenetic analysis of the corresponding sequences but there are no established standards on how closely related STs should be to be grouped together under the same CC (1). Rather, CCs are usually defined by the clustering patterns (e.g., monophyly) of STs, and after overlaying epidemiological data, that is, in an empirical way. If the intra-species diversity was organized in consistent, discrete units across different taxa, this would have provided for more natural and precise intra-species unit definition(s) compared to the existing practice and thus, improved communication about intra-species diversity. While it has been recently recognized that prokaryotic organisms may form such discrete units at the species level (6), it remains unclear whether or not such consistent units across different taxa also exist within species.

Specifically, culture-independent (metagenomic) studies of natural microbial populations during the past decade revealed that bacteria and archaea predominantly form sequence-discrete populations with intra-population genomic sequence relatedness typically ranging from ∼95% to ∼100% ANI depending on the population considered. For example, younger populations since the last population diversity sweep event show lower levels of intra-population diversity. In contrast, ANI values between distinct populations are typically lower than 90% (7). Intermediate identity genotypes, for example, sharing 85–95% ANI, when present, are generally ecologically differentiated and scarcer in abundance, and thus should probably be considered distinct species (6, 8, 9) rather than representing cultivation or other sampling biases (10). Such sequence-discrete populations have been recovered from many different habitats, including marine, freshwater, soils, human gut, and biofilms, and are usually persistent over time and space [e.g., (11–15)] indicating that they are not ephemeral but long-lived entities. Further, these sequence-discrete populations commonly harbor substantial intra-population gene content diversity (i.e., they are rarely clonal) (11, 14). Therefore, these populations appear to be “species-like” and may constitute important units of microbial communities. Moreover, the 95% ANI threshold appears to be largely consistent with how isolate genomes have been classified into (named) species in the last couple decades; that is, ∼97% of named species include only organisms with genomes sharing >95% ANI (16). In summary, it appears that a natural gap in ANI values can be used to define prokaryotic species and has been largely consistent with how species are recognized (16). In this context, “discontinuity” or “gap” refers to the dearth of genome pairs showing 85–95% ANI relative to counts of pairs showing ANI >95% or <85%. Whether or not a similar ANI gap exists within species has not been evaluated yet.

A related term that is also commonly used to catalogue intra-species diversity is strain. The “strain” represents a fundamental unit of microbial diversity that is commonly used across medical or environment studies to designate the smallest distinguishable unit, presumably a unit within STs. Unlike the stringent and precise definition of ST and CC, the definition of a strain is more relaxed and often appears context dependent. Specifically, the concept of a strain is primarily based on the notion of pure cultures (17). The Bacteriological Code defines strain as the group “*of the descendants of a single isolation in pure culture*” (18). Accordingly, a strain is expected to represent a genome or a collection of genomes that have no single-nucleotide or gene content differences or, if such differences exist, they are expected to not encode for important phenotypic differences (19). Unfortunately, this definition of a strain – while being commonly used across microbiological fields – remains problematic, and often leads to confusion in communication. Notably, it is not always clear when two distinct genomes or cells should be considered the same or separate strains since cell ancestry information is often missing, such as in environmental (culture-independent) surveys. Additionally, the isolation of an organism (wild-type) in the laboratory is frequently accompanied by phenotypic changes due to adaptation to laboratory conditions; yet, the wild-type and the lab-adapted cells are typically considered the same strain (17), even though their observed phenotypes may not fully overlap in some cases. The existence of phenotypic differences among members of the same strain could be confusing because high phenotypic similarity is expected at this (the strain) level. To circumvent some of these limitations, we have recently proposed a definition threshold for strains at 99.99% ANI on the basis of high gene-content conservation (>99.0% of total genes, typically) and thus, phenotypic relatedness among 138 isolate genomes of *Salinibacter ruber* recovered from two saltern sites in Spain (20). However, testing this definition with a larger collection of genomes and how it precisely relates to STs and CCs have not been performed yet.

## Results and Discussion

### An ANI gap within species around 99.2-99.8%

In the process of assessing cultivation biases as a possible explanation for the ANI-based sequence discrete populations previously (6), we observed another discontinuity (or gap) in ANI values that may be used to more reliably and systematically define the units within a species. Specifically, the analysis of 18,123 complete genomes from 330 species available in NCBI’s Assembly database with at least ten genome representatives per species revealed a clear bimodal distribution in the ANI values within named species or 95% ANI-defined groups of genomes (genomospecies). That is, there is a scarcity of genome pairs showing 99.2-99.8% ANI (average around 99.5% ANI) in contrast to genome pairs showing ANI >99.8% or <99.2%. Specifically, among the 18,123 complete genomes in our dataset, there are 4,280,133 genome pairs showing ANI >96%, which would translate to about 107,000 pairs per every 0.1 percent unit of ANI if there was no bimodal distribution and the ANI values among these genome pairs were evenly distributed between 96% and 100% ANI. Our analysis revealed only 235,527 genome pairs between 99.2% and 99.8% ANI, which is three-fold fewer data points than expected by chance alone in a uniform ANI value distribution (642,000 pairs expected). No other ANI range within 96-100% had such a strong bias based on our dataset. That is, a pronounced gap in ANI values is observed among very closely related members of a species around 99.2-99.8% ANI (Fig. 1). Importantly, this ANI gap appears to be consistent across phylogenetically diverse species from a dozen of distinct bacterial phyla evaluated, including gram-negative and gram-positive. About half of the 330 species evaluated appear to be environmental species (as opposed to pathogen or human/animal host-associated) or species of biotechnological interest, revealing no major bias based on the (presumed) ecology of the organisms evaluated (Fig. S1). Further, the ANI gap does not seem to be driven by a couple or a few species based on a sub-sampling of all species to the same number of genomes (n=10) (Fig. S2). Instead, it represents a nearly universal property of the 330 species evaluated (see also Fig. S3 for specific species examples). Therefore, it appears that another important level of genomic differentiation may exist within species.

**Figure 1.**
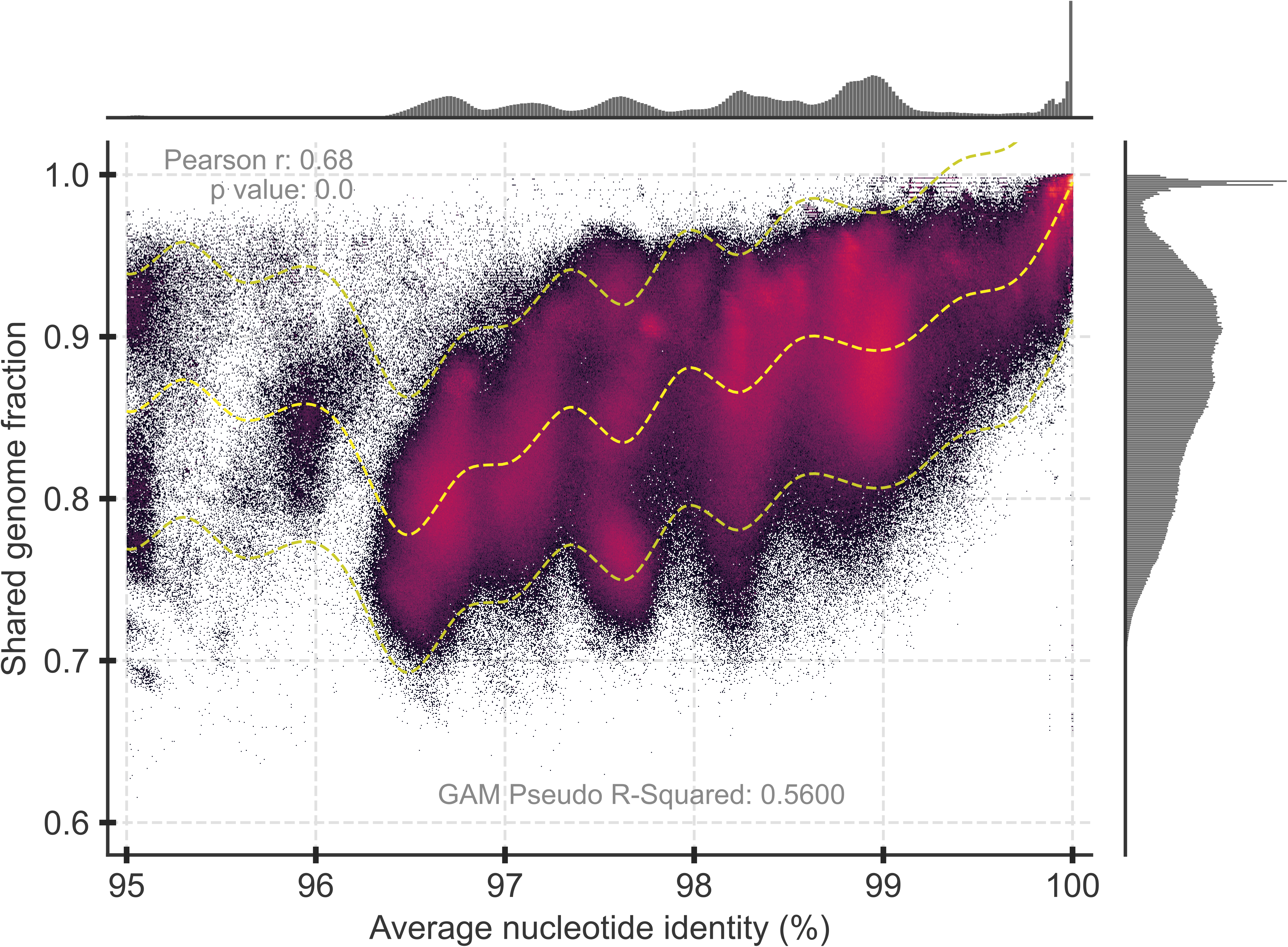
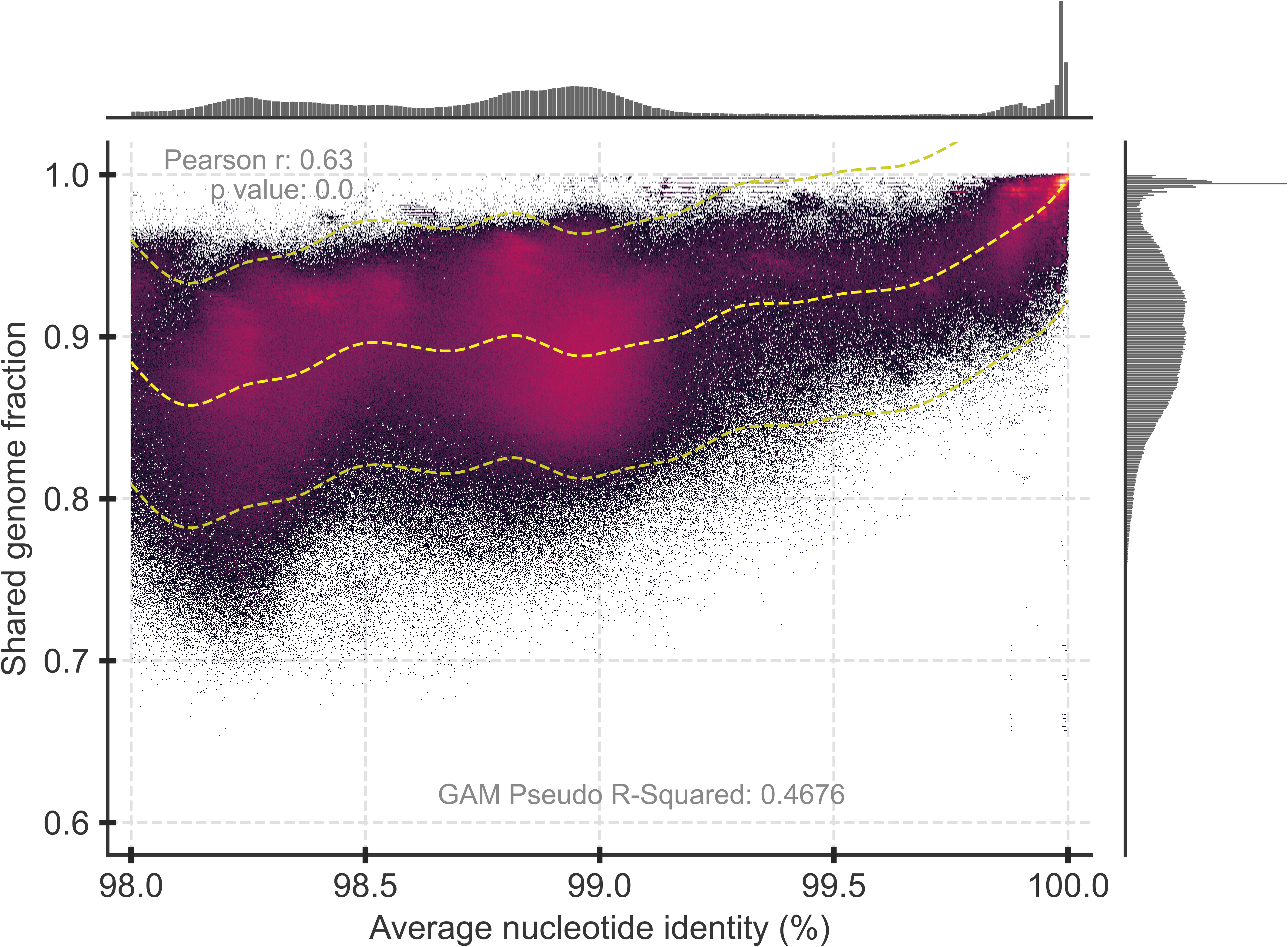
ANI vs. shared gene content for the 17,283 complete genomes used in this study. Each datapoint represents a comparison between a pair of genomes. FastANI (16) was used to generate ANI values between the genomes of a pair (x-axis) and their shared genome fraction (y-axis). The shared genome fraction was calculated by dividing the number of bidirectional fragment mappings over the total query fragments determined by FastANI. Only a single set of values is reported per pair, the one that used the longer genome as the reference (and the reverse comparison was omitted). Note that only datapoints representing genome pairs sharing ANI >95% are shown (n=4346079 datapoints), and that panel B is a zoomed-in version of panel A (n=2676181 datapoints). The main scatter plot is shaded by the density of points using the Datashader package in Python with Matplotlib. The trendline was calculated using linearGAM from pyGAM and includes the 95% confidence interval. The marginal plots outside the two axes show histograms for the density of datapoints of each axis. Note the low-density region in the ANI value distribution around 99.2-99.8% (Panel A), which becomes more obvious when zooming in to the 98-100% ANI range (Panel B).

It is unlikely that this 99.2-99.8% intra-species ANI gap is due to cultivation or classification biases due the reasons mentioned previously, such as that cultivation media usually do not distinguish between members of the same or closely related species (i.e., similar organisms) (6), and that random subsampling provided similar patterns (Fig. S2). It is also highly likely that the intra-species ANI gap is even more pronounced in nature because very closely related genomes (e.g., showing ANI >99.8% to each other) are often selected against for genome sequencing (and thus, are likely underrepresented in our collection) based on pre-screening using fingerprinting techniques (e.g., RAPD, MLST) in order to avoid sequencing of redundant genomes. Further, we were not able to identify clear exceptions to this 99.2-99.8% intra-species ANI gap when examining individual species with enough sequenced representatives, although such exceptions likely exist. For instance, several species in our collection did not have enough highly related genome representatives (showing ANI >99% to each other) to assess the critical area of ANI value distribution (i.e., the 99-100% range), and this could be due to the pre-screening biases mentioned above or reflect their actual natural diversity patterns (see also Fig. 4, Panel D for an example of an exception within *E. coli*). Further, for a few species (n < 10) such as *Listeria monocytogenes* and *Bordetella bronchiseptica*, the intra-species ANI gap appears to exist but is shifted compared to the 99.2- 99.8% ANI that characterized most well-sampled species (Fig. S3; all 330 species are available on the GitHub at https://github.com/rotheconrad/bacterial_strain_definition). Therefore, for future studies, we suggest evaluating the ANI value distribution for the species of interest, and if the data indicate so, to adjust the ANI threshold to match the gap in the observed ANI value distribution. The 99.2-99.8% ANI range should represent the gap for most species based on the dataset evaluated here.

### Support from a high-throughput cultivation environmental study and long-read metagenomes

Our team has recently described a collection of high-draft, isolate genomes of *Salinibacter ruber* (n=162), chosen for sequencing at random from a larger collection of isolates (n=257) recovered from two solar saltern sites on the Mallorca and Fuerteventura Islands (Spain) (9). Solar salterns are human-controlled environments used for salt production. They are operated in repeated cycles of feeding with natural saltwater, followed by water evaporation due to ambient sunlight, and finally, salt precipitation. Salterns from different parts of the world have been reported to harbour similar microbial communities, generally consisting of two major lineages, i.e., the archaeal *Halobacteria* class and the bacterial family of *Salinibacteraceae*, class *Rhodothermia* (21). Within each lineage, the species richness is relatively high (22–24). *S. ruber* often makes up 5-20% of the total microbial community of saltern sites, i.e., it is an abundant population *in situ*, and the growth media used in our previous study have been tested and found to not bias the diversity of *S. ruber* isolates that can be recovered in culture (9, 21). Notably, our analysis of the 162 genomes revealed that only 0.35% of the total 13,122 ANI comparisons fell between 99.6% and 99.8% vs. 1.24% expected if the total ANI values higher than 99% were distributed uniformly at random between 99% and 100% (∼4-fold reduction in datapoints; Fig. 2). Therefore, the intra-species ANI values between members of the same *S. ruber* species from a single sampling site and year (e.g., Mallorca Island) or two sites separated by about 2,000 km (Mallorca vs. Fuerteventura Islands) revealed a remarkably similar ANI gap to that observed with the heterogenous genome collections available in NCBI.

**Figure 2.**
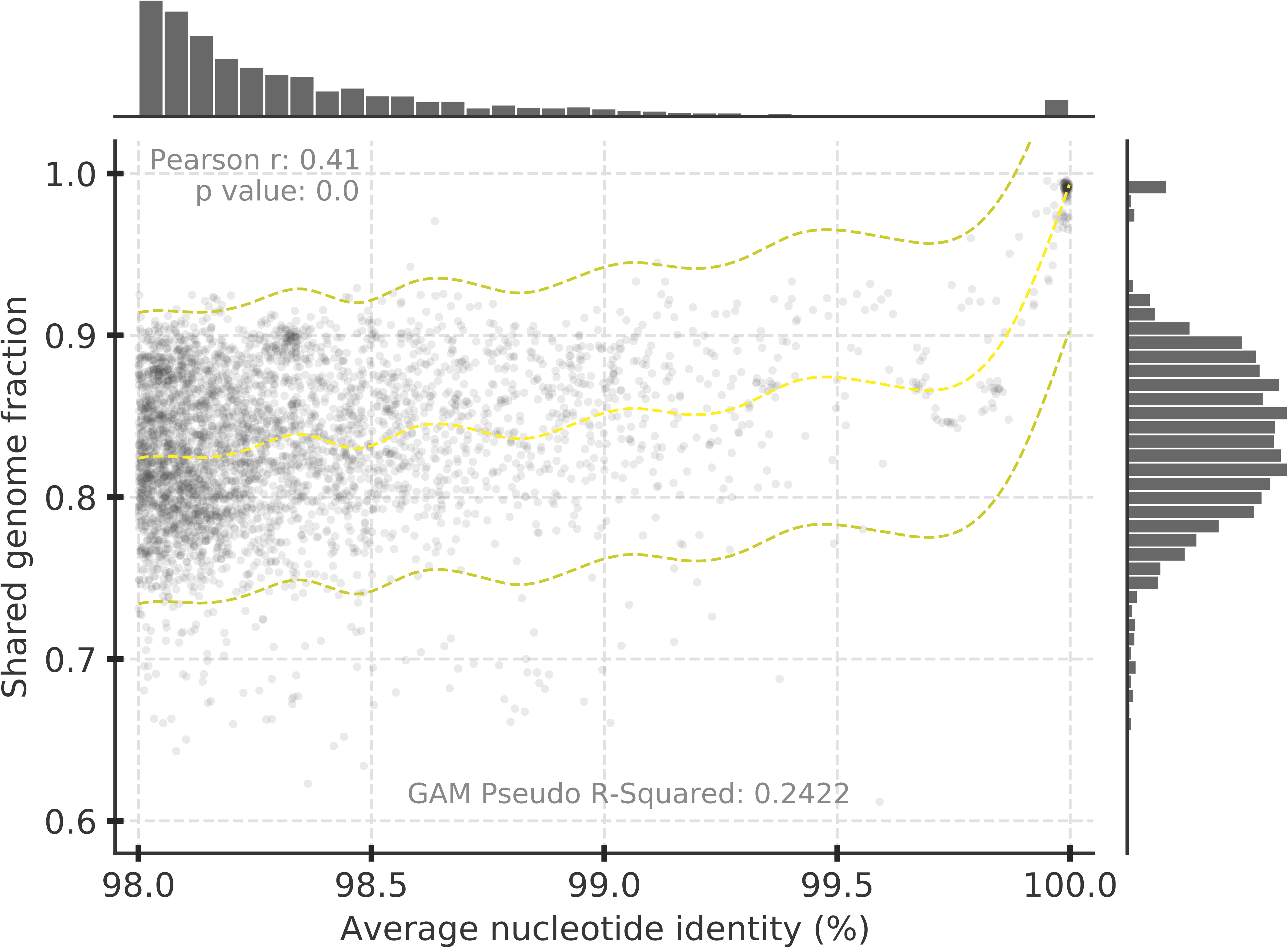
ANI vs. shared gene content for *Salinibacter ruber* isolate genomes recovered for the same site. Fig. 2 is identical to Fig. 1B except that the underlying data represent 162 *S. ruber* isolate genomes (n= 3847 datapoints with >98% ANI) recovered from two saltern sites on the Mallorca and Fuerteventura Islands (Spain) as previously described (9). Note the lack of genome pairs showing ANI values between 99.2% and 99.8% and the sharp increase in gene-content differences in pairs of genomes showing ANI <99.8%, which echoed the results obtained with NCBI genomes (Fig. 1). Similar results were obtained when the genomes from each site were analyzed independently (data not shown).

We also examined recently available long-read metagenomes from a variety of habitats (25, 26) to offer a culture-independent assessment of the intra-species diversity patterns. It should be noted that short-read metagenomes are not ideal for our purposes because the shorter sequence fragments show a larger dispersion of identity values around the (genome-) average identity value and thus, could mask (obscure) the ANI gap we observed in the analysis of both NCBI complete and *S. ruber* draft isolate genomes described above. Moreover, direct read mapping of short reads can only offer identity resolutions of 1 divided by the read length (e.g., 1% resolution for 100 bp, or 0.4% resolution for 250 bp), making it impossible to resolve a gap so close to 100% identity. Nonetheless, previous short-read metagenomic surveys that recovered and compared metagenome-assembled genomes (MAGs) of the same species from different samples have revealed a scarcity of MAGs that are related around 99.0-99.5% ANI (15), although the resolution – and thus, the sharpness of this ANI gap – is not as great as that presented here based on isolates above. The latter could be due, at least in part, to the assembly process frequently merging sequences related at 97-98% nucleotide identity or higher into a consensus sequence.

To avoid the possibility of the assembly obfuscating sequence similarities in the data, we did not assemble the long-read sequences but made direct comparisons against each other or against a reference genome or MAG recovered from the same sample, using only reads longer than 10 Kbp and from both Oxford Nanopore and PacBio datasets. We also examined, independently, the diversity patterns based on full sequences of the ribosomal polymerase subunit B (*rpoB*) gene recovered by a subset of the long-reads to offer higher resolution, since *rpoB* has been shown to represent a reliable marker that reflects well the whole-genome identity (5). The analysis of *rpoB* metagenomic sequences also circumvented the effect of varied degrees of sequence conservation between different genes on the signature; i.e., that some genes are more or less conserved than the genome average, which could introduce noise with respect to gaps in nucleotide diversity patterns. Collectively, the results showed that the intraspecies ANI gap is present in most populations that were abundant enough to be adequately sampled by the metagenomic library in the human gut (Fig. 3) as well as soil and ocean habitats (Fig. S4). In a few cases, especially of oceanic populations, the ANI gap was not observed, mostly because either the populations were too clonal to assess (e.g., no reads showed nucleotide identities <99%; majority of the cases observed) or harbored extensive intra-population that was – more or less – evenly distributed between 96% and 100% ANI (Fig. S4, B and C). For several of the latter cases, however, the ANI gap became (more) obvious when the analysis was restricted to the *rpoB* gene (Fig. S4D), revealing that the effect of varied degrees of sequence conservation between different genes on (blurring) the signature was significant even based on long reads. Collectively, these results showed that while exceptions to the pattern exist, the predominant picture is that the signature is present in natural populations, providing further support for the intra-species ANI gap revealed by the analysis of the isolate genomes described above.

**Figure 3.**
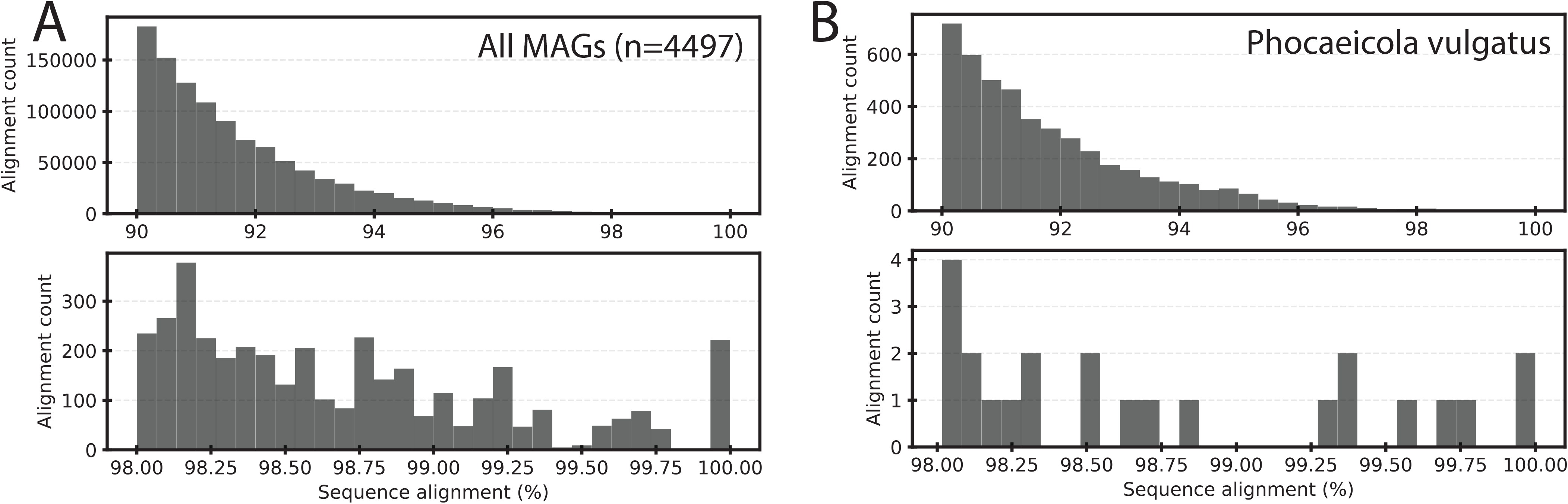
The intra-species ANI gap is present in natural populations recovered by long read metagenomic sequencing of human fecal samples. The underlying data are Oxford Nanopore long read metagenomes of human fecal samples (109 samples, about 136Gbp, in total, after read quality trimming) described previously (26). The graphs show the nucleotide (nt) identity distribution of individual long reads against the MAGs recovered by the same study. All reads were quality-trimmed with Filtlong (v0.2.1) for minimum read lengths of 1kb and average quality scores of >90% across 1kb sliding windows. Reads were mapped non-competitively with minimap2 (v2.24) allowing for spliced alignments (-x splice). Only alignments sharing at least 20% sequence overlap with a MAG were reported. Similar distributions were obtained when reads were simply mapped all versus all (data not shown). Top histograms show all reads sharing nt identities above 90%; bottom histograms show the subset of these reads that share nt identities above 98%. Panel A summarizes the mapping of all long reads against all MAGs (n=4,497). Panel B shows mapping of all long reads against a single MAG taxonomically identified as *Phocaeicola vulgatus*. Note the sparsity of reads mapping between 99.4-99.9% nt identity for panel A and 98.8-99.3% for panel B.

### Gene content diversity within the >99.8% ANI clusters

Another notable observation from the data from all species comparisons is that shared gene content generally decreases as ANI distance (or genomic divergence) increases within the 95% ANI clusters, but the decrease is biphasic. That is, shared gene content decreases quickly among genome pairs sharing 99.8-100% ANI but then, the decrease is less dramatic in genome pairs sharing between 96.0%-99.8% ANI. In other words, genome pairs sharing between 99.8%- 100% ANI (i.e., one fourth of a unit of ANI) may differ in their total gene content by up to 10% (average values of genome pairs showing ∼99.8% ANI) and more divergent genomes of the same species (i.e., showing 96.0%<ANI<99.8%) may differ by up to 20% (average values of genome pairs showing ∼96.0% ANI), adding another ∼10% of gene content differences for 3.75 additional units of ANI (vs. 0.25 unit in the 99.8-100% range). Further, the genes that differed between genomes sharing >99.8% ANI are more enriched in hypothetical and mobile (e.g., prophage and transposases) functions compared to the functions that differ between more divergent genomes by ∼10% of the total genes in the genome (p<0.001, z-test; Fig. S5), consistent with what was reported earlier for intra-species gene-content diversity (27). Collectively, these results show that genome pairs showing ANI >99.8% are also expected to be much more similar in gene-content compared to more divergent genomes of the same species. Nonetheless, it is important to highlight that even very closely related genomes (showing ANI >99.8%) often show substantial gene-content differences, up to about 10% of the total genes based on our evaluation, albeit most of these differences are likely ephemeral and metabolically/ecologically not-important genes (e.g., Fig. 5S). Hence, members of the same 99.8%-ANI-based intra-species unit should be expected to be overall more similar in shared functional gene content and thus, phenotype relative to comparison between such intra-species units, but not necessarily identical.

### Comparison to Sequence Types (or STs)

We also assessed how consistent the 99.5% ANI threshold (as the midpoint of the 99.2- 99.8% range that corresponds to the gap) is with the assignment of genomes to other intra-species units. We found that the 99.5% ANI threshold is most similar to the Sequence Type (ST), the latter defined as identical sequences for 6-7 genetic loci, among the units evaluated (data not shown). We primarily report on the analysis of the *E. coli* species below because it is a good representative of the ANI patterns observed within other species in our dataset, the large number of closed *E. coli* genomes available (n=1218), and the availability of a robust Multi- Locus Sequence Typing/Analysis (MLST/MLSA) scheme (28) that has been used for at least two decades to provide below-species resolution and identify outbreaks of *E. coli* pathogens. Under the *E. coli* MLST scheme, genomes are assigned to the same ST based on identical sequences in seven *E. coli* core genes (namely, *adk*, *fumC*, *gyrB*, *icd*, *mdh*, *purA*, and *recA*) (28). Our evaluation showed that the 99.5% ANI threshold is largely consistent with how genomes are assigned to STs; that is, ∼80% of ST assignments, for the four most abundant STs (n=615 genomes), were supported by the 99.5% ANI threshold (Fig. 4). In other words, only about 20% of the genomes that were assigned to the same ST showed <99.5% ANI among themselves (high recall). The existence of a clear gap around 99.5% ANI in the latter cases indicated that these STs could be split in two (or more) STs for more homogenous STs in terms of overall genomic relatedness (e.g., ST-10 and ST131, Fig. 4). The higher resolution provided by ANI in these cases is due, at least in part, to the fact that the core genes used in current MLST schemes tend to be more highly conserved, at the sequence level, than the genome average (represented by ANI). These results are also consistent with our previous conclusion that the 95% ANI threshold for species demarcation should be adjusted upwards if the ANI is based on a few universal or core genes, as opposed to the whole-genome (16). Further, ∼3.5% of the genomes with ANI >99.5% were assigned to different ST, revealing even higher precision (Table S1; Suppl. Fig. S6). Similar results to those reported here for *E. coli* were observed for several of the 14 most-sampled species in our dataset with available MLST schemas, albeit recall was slightly better than that observed for the *E. coli* dataset (average across all ST comparisons: 83.1%, stdev 0.31) while precision was worse (average 91.6%, stdev 0.19).

**Figure 4.**
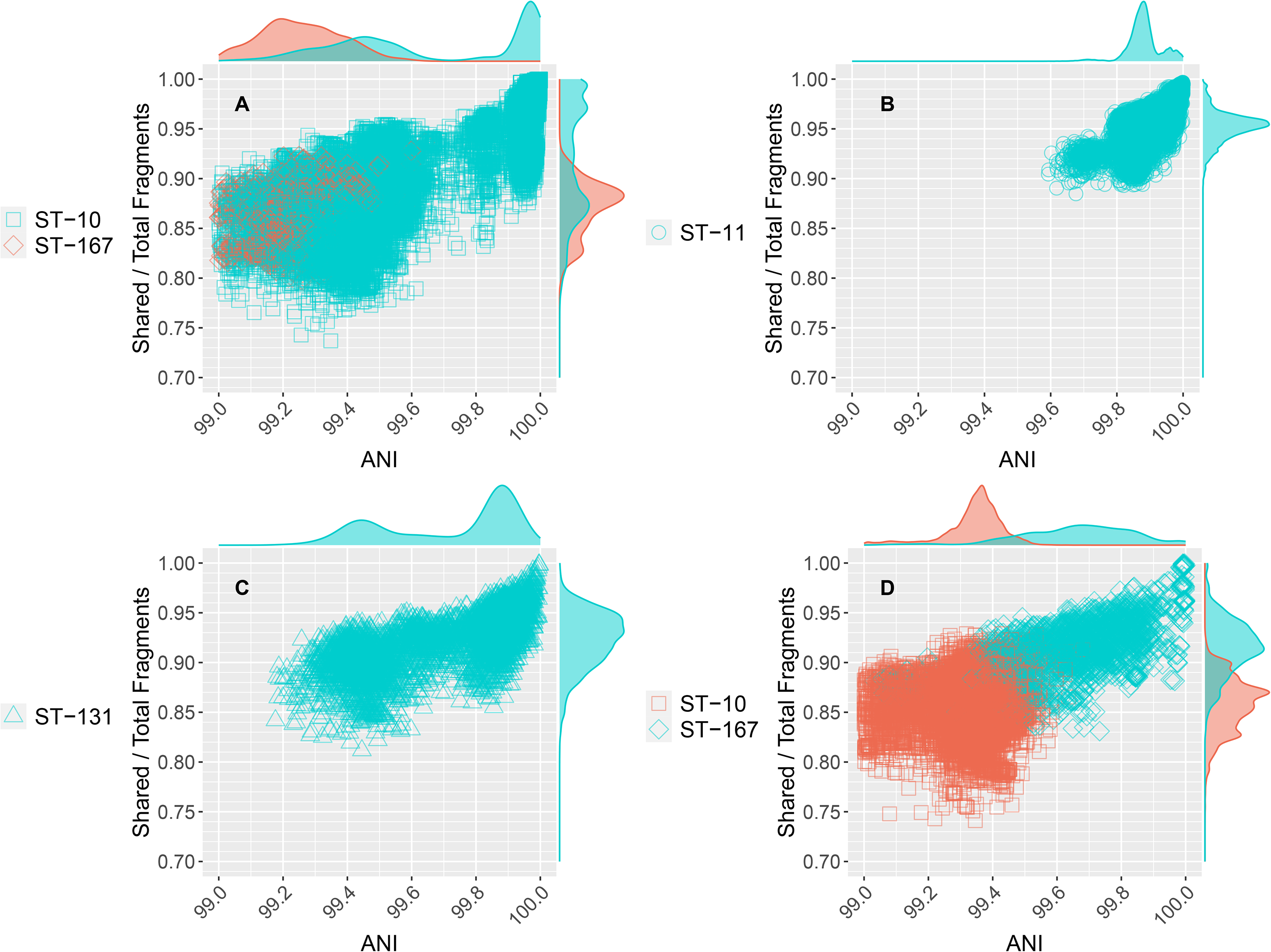
Comparison of the 99.5% ANI threshold to available Sequence Types (STs) of *E. coli*. All *E. coli* genomes were assigned to a ST using the tool mlst version 2.19.0 (28). The four panels show the four most abundant STs based on the number of genomes assigned to them (see Table S1 and Fig. S6 for underlying data and F1 statistic, respectively). Each datapoint is a comparison between two genomes, similar to those shown in Fig. 1; the marginal plots show the kernel density estimate of datapoints for each axis. Datapoints are cyan if both genomes in the pair were assigned to the same, reference ST; red datapoints represent pairs for which one of the genomes in the pair is assigned to a closely related, yet distinct ST than the reference ST. Note that for ST-11, recall and precision of 99.5% ANI vs. ST designation is perfect because there are no genomes, and thus STs, that are closely related to ST-11, which is also consistent with a pronounced ANI gap at 99.5% for ST-11, and the substantial overlap in terms of ANI values between the closely related ST-10 and ST167 (low recall). ST-131 and ST-10 appear to harbor too much genomic diversity and could be split in more than one ST based on the 99.5% ANI criterion (and the bimodal ANI value distribution around 99.7% ANI). ST-167 represents a possible deviation from the main pattern, meaning no clear ANI gap is obvious for this ST around 99.5% but a gap may be obvious at lower ANI values, around 99.0%-99.2%. A couple more similar examples to ST-167, out of about 330 cases examined, were noted (Fig. S3 and https://github.com/rotheconrad/bacterial_strain_definition), indicating that exceptions to the main pattern are generally infrequent.

The high precision (and recall) of the 99.5% ANI threshold compared to ST assignments is due, at least in part, to the fact that hundreds, if not thousands (e.g., at least 3,000 for the *E. coli* pairs), of genes are used in each ANI calculation at this level of high relatedness. Thus, the ANI threshold is robust against horizontal gene transfer or other evolutionary events that could affect sequence identity of one or a few loci, a known limitation of traditional MLST approaches. Importantly, the 99.5% ANI threshold, or a different ANI threshold to better capture intra-species diversity patterns as recommended above, is a property that emerges from the data themselves as opposed to a manmade threshold (such as “identical sequences in seven loci” used in MLST applications) and thus, it should capture the natural diversity patterns better. Consistent with this interpretation, our preliminary results from applying the 99.5% ANI threshold to a collection of *E. coli* isolate genomes collected over a period of 18 months in Northern coastal Ecuador as part of the EcoZUR study (for “*E. coli* en Zonas Urbanas y Rurales”) (29) shows that the 99.5%-ANI-defined STs map well to local outbreaks of pathogenic *E. coli* (Feistel et al., in preparation). Therefore, using the 99.5% ANI to define new or refine existing STs could provide data-informed groups that encompass genomically more homogenous organisms compared to the existing practice in some cases. Another important advantage of the 99.5% ANI is that it can be automatically implemented and thus, does not require manual curation, which is the case when establishing new ST numbers when novel (meaning, not seen previously) sequences become available (3).

### ANI-based definitions for strain, sequence type (ST) and clonal complex (CC)

Our evaluation shows that the 99.5% ANI threshold, or the 99.8% if the upper end of the observed ANI gap is used instead, is directly comparable to how STs have been defined (e.g., Fig. 4) and thus, could be used to complement or even substitute in the fugure the requirement for 6-7 identical loci that has the shortcomings explained above. We propose the midpoint (99.5% ANI) as opposed to the upper value (99.8% ANI) of the gap as a more conservative threshold and in order to account for the variation observed among different species with respect to the ANI value range that corresponded to their own gap (e.g., Fig. S3). We also suggest that the term genomovar could be used to refer to these 99.5%-ANI intra-species units, instead of STs, in case that ST should maintain its original conception of 6-7 identical loci for historic, pragmatic or other reasons. The term genomovar was originally used to name distinct genomic groups within species that cannot be distinguished phenotypically from each other and therefore, cannot be classified as distinct species based on the standard taxonomic practices (30). Hence, genomovar may capture conceptually well the 99.5% ANI groups, as we also proposed recently elsewhere (20).

We think that CCs should continue to be defined based on phylogenetic analysis of STs; in fact, we see the 99.5% ANI threshold proposed here as a complementary, rather than competitive, approach to the phylogenetics because it provides convenient means to define and/or refine STs, which can then be used in phylogenetic analysis to define CCs, assess evolutionary relationships between STs, etc. Related to this, it is important to note that the computation of ANI is, on average, two orders of magnitude faster compared to the phylogenetic placement of a genome using all core genes (16). Therefore, the ANI-based approach and thresholds proposed here should provide highly efficient and practical means to classify (type) large collections of genomes into STs at the whole-genome level and subsequently perform phylogenetic (or other) analysis of representatives of the resulting STs.

Finally, we did not observe another ANI gap within the 99.5% ANI clusters and thus, recommend the use of the term strain only for nearly-identical genomes. We recently proposed to define a strain as a collection of genomes sharing ANI >99.99% based on the extent of gene-content differences among the *S. ruber* genomes originating from the same saltern site (20). This threshold ensures high gene content similarly, e.g., typically, >99.0% gene content is shared at this ANI level based on the *S. ruber* genomes (20) and the data presented here (Fig. 1 and Fig. 2). Further, our previous study showed that this ANI level encompasses well the typical sequencing and assembly noise observed when splitting the raw reads from a *S. ruber* isolate genome project in two halves and assembling the two subsets independently for comparisons of the resulting genome sequences (20). It should be noted, however, that 99.99% ANI threshold represents only practical and convenient means for defining strains, and it could be neglected or adjusted should key phenotypic differences distinguishing organisms sharing ANI >99.99% are known/found such as antibiotic resistance or catabolic genes carried by plasmids.

### What are the underlying mechanisms for the 99.5% ANI gap?

The mechanism(s) that underly the 99.5% ANI gap (or the earlier 95% ANI gap for the species level) remain essentially speculative and should be the subject of future research in order to further advance the mechanistic understanding of the microbial diversity patterns observed in nature. Most notable is the idea that members of a population cohere together via means of unbiased (random) genetic exchange which is more frequent within vs. between populations or CCs (i.e., *the biological* or *sexual species concept*) (31). A competing hypothesis is that several members of the species are functionally differentiated from each other either due to specialization for different growth conditions or different affinities for the same energy substrate and thus, selection over time for these functions purge diversity (i.e., *the ecological species concept*) (32–34). It is intriguing to note that the ecological explanation is also consistent with the notion that STs or different strains of the same species are somewhat ecologically and/or functionally distinguishable from each other. Notably, given an estimated mutation rate of ∼4x10^-10^ per nucleotide per generation (35) and between 100 to 300 generations per year (36), it would take two distinct *E. coli* lineages or STs at least twenty thousand years since their last common ancestor to accumulate 0.5% difference (i.e., fixed mutations) in their core genes or 99.5% ANI. Therefore, there is enough time, at least theoretically, for the ecological purging of diversity to take place at around the 99.5% ANI level and thus, account for the ANI patterns observed herein. Intriguingly, it has been shown that the explicit inclusion of extinction events in a neutral model of evolution can also result in punctuated distributions of genetic differentiation, opening up a third possibility of historical contingency from stochastic events (37). However, we note that while stochasticity can explain bimodal (or multimodal) distance distributions, a scarcity of ANI values in the exact same range (i.e., around 99.5% ANI) would be unlikely to repeatedly emerge by chance alone across many different species with distinct lifestyles and evolutionary tempo, as opposed to this range varying between species. In any case, the data available in support of one of these (or another) hypotheses remain sparse and/or anecdotal to date, to the best of our knowledge, and the analysis presented in this study did not aim to advance this issue further but rather to present a highly intriguing observation of patterns of diversity that could have major practical consequences in defining strains, STs, and species more broadly.

## Conclusions

Regardless of what the underlying mechanisms are for the 99.5% ANI gap, the results presented here show that the patterns of natural diversity among thousands of sequenced genomes are consistent with a 99.5% ANI threshold that can be used to identify STs more reliably and precisely compared to the current practice. Regarding the use of this threshold to define (or refine) STs vs. strains, we believe that the threshold is highly appropriate, as well as it matches well the intended meaning and use of STs, and thus its application to ST definition is straightforward, although we recommend the alternative term of genomovar to avoid confusion with historical ST definitions. Notably, recent large-scale surveys of the human gut microbiome (38) have also revealed distinct intra-species units closely matching the 99.5% ANI threshold and genomovar definition proposed here, even though these units were often called strains previously (39). However, for the strain level, genomes showing ∼0.5% or ∼0.2% difference in ANI (99.5% or 99.8% ANI, respectively) often show substantial sequence and gene content divergence (e.g., Fig. 1) and thus, phenotypic differences. Therefore, multiple (distinct) strains should be expected to be grouped together within the same 99.5% ANI cluster in such cases, and strain, in general, represents a more fine-grained level of resolution than the 99.5% ANI level. The results presented here based on all well-sampled bacterial species further reinforced our recent proposal based on the *S. ruber* isolate genomes from the same site to use 99.99% ANI as the threshold to define strains (20). Collectively, we expect that the findings reported here will advance the molecular toolbox for accurately delineating and following the important units of diversity withing prokaryotic species and thus, would greatly facilitate future epidemiological and micro-diversity studies.

## Methods

Step by step methods, including how average trendlines were fit to the data, custom Python code, NCBI Assembly accession numbers for selected genomes, and plots for each selected species are available from: https://github.com/rotheconrad/bacterial_strain_definition. Briefly, all genome sequences were obtained from NCBI’s RefSeq Assembly database on April 20^th^, 2022 and were labeled as “complete” and “latest”. ANI values and the shared genome fractions were directly obtained from the output of FastANI version 1.32, “One to Many” mode with default settings (16). Results were concatenated to create within species all vs. all output. Self matches were removed, and genome pairs were filtered by minimum ANI values according to the axes of each figure. Selected individual species plots (i.e., all vs. all output) of shared genome fraction vs. ANI are shown in the Supplementary Material. *E. coli* genomes were assigned to sequence types (ST) using the using the command-line tool mlst (https://github.com/tseemann/mlst) version 2.19.0 (28) with default settings.

## Acknowledgments

This work has been supported by the US National Science Foundation (Award No 1759831 and 2129823) to KTK. LO and RA acknowledge funding by the Max Planck Society.

## Code and data availability

All code and data details are available from https://github.com/rotheconrad/bacterial_strain_definition.

## Competing interests

The authors declare no competing interests.

## Author contact list

Luis M. Rodriguez-R

Roth E. Conrad

Dorian J. Feistel

Tomeu Viver

Fanus Venter

Luis Orellana

Rudolph Amann

Ramon Rossello-Mora

Konstantinos T. Konstantinidis

## Notes

https://github.com/rotheconrad/bacterial_strain_definition

### Competing Interest Statement

The authors have declared no competing interest.

### Summary of Updates

General wording changes to clarify communication

