## Supplemental Material for "An ANI gap within bacterial species that advances the definitions of intra-species units"

*equal first authors.

**Keywords:** ANI, strain definition, micro-diversity, epidemiology, clonal complex

**SUPPLEMENTARY FIGURES AND TABLES**.

**Supplementary Table S1. Statistics of ST separation at different thresholds of ANI**. The first header row indicates the different thresholds evaluated to separate STs. True Positives (TP) were defined as genome pairs with ANI > threshold and the genomes were assigned to the same ST; True Negatives (TN) when the genome pair had ANI < threshold and the genomes were assigned to different STs; False Positives (FP) when the genome pair had ANI > threshold and genomes were assigned to different STs; and False Negatives (FN) when the genome pair had ANI < threshold and genomes were assigned to the same ST. Precision, Recall, Accuracy, and F1 score were defined as shown on the Table (bottom).

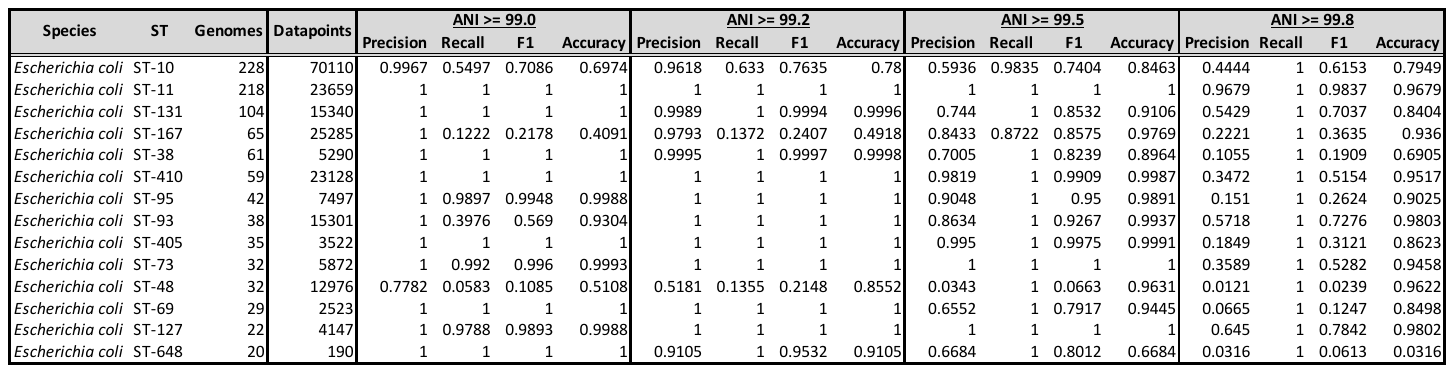

Precision = TP / (TP + FP)

Recall = TP / (TP + FN)

F1 score = (2 * Precision * Recall) / (Precision + Recall)

Accuracy = (TP + TNP) / (TP + TN + FP + FN)

**Supplementary Figure S1. Clinical and environmental genomes show similar intra-species ANI value distribution patterns**. The Fig. S1 is identical to Fig. 1 except that the 17,283 NCBI genomes were separated in **clinical** (n =  224 species; 27150 genomes; 2585561 genomes pairs with >95% ANI), for those associated with human or animal hosts (top), and **environmental** (n =  113 species; 3706 genomes; 61212 genomes pairs with >95% ANI), for those associated with biotechnological applications and/or no-host (bottom), based on their NCBI record. Note that while there are fewer environmental genomes overall, these do show the ANI gap around 99.5%.

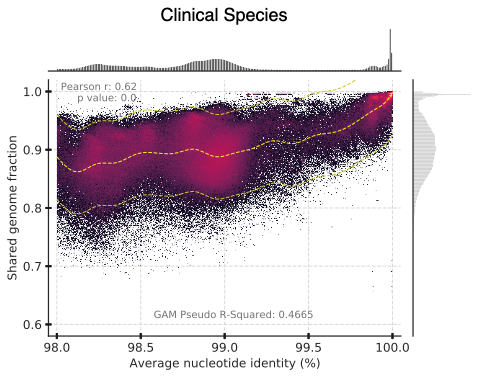

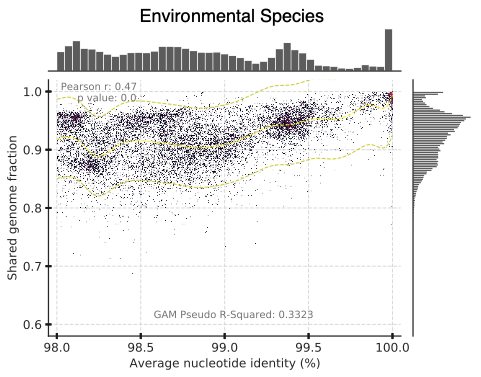

**Supplementary Figure S2. Random subsampling of the genomes supports the 99.5% ANI gap.** The figure is similar to Fig. 1 but only includes 256 (top Panel, 256000 genome pairs with >95% ANI) and 215 (bottom Panel, 215000 genome pairs with >98% ANI) species with exactly 45 datapoints (genome pairs; from all vs. all comparisons of 10 genomes) each. Note that the 99.5% ANI gap is apparent, albeit not as pronounced as in Fig. 1 due to the subsampling of the datapoints.

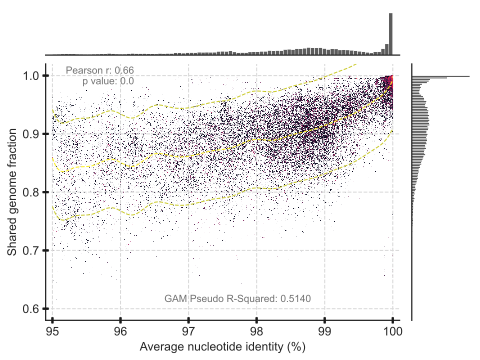

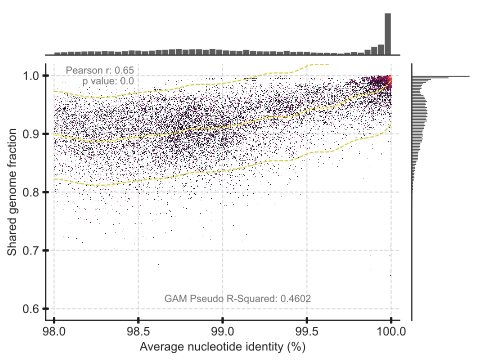

**Supplementary Figure S3. Examples of individual species showing the 99.5% ANI gap (top) and species that deviate from this pattern (bottom).** The figure is similar to Fig. 1 but only shows datapoints from individual species, one species per panel (see panel title).

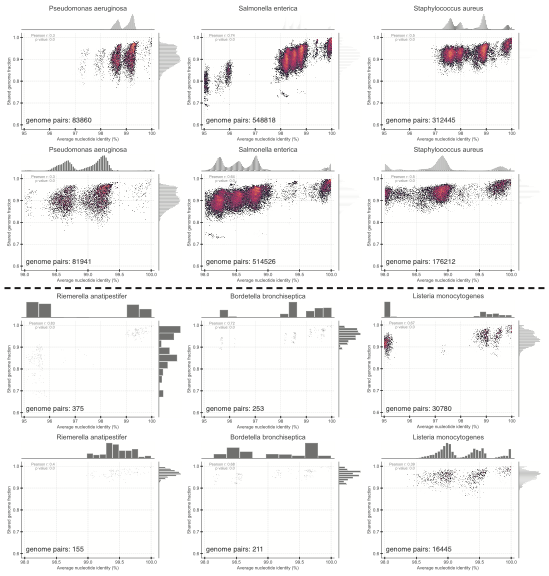

**Supplementary Figure S4. The intra-species ANI gap is often present in natural populations recovered by long-read PacBio metagenomic sequencing from the surface waters of the North Sea**. The underlying dataset is a PacBio HiFi long-read metagenome from surface waters of the North Sea after a seasonal algal bloom described previously (1). The graphs show the nucleotide (nt) identity distribution of individual reads against a reference MAG recovered from the same dataset. All reads shared at least 5Kbp of their >10Kbp length with the MAG for stringent results while similar distributions were obtained when reads were searched against themselves (data not shown). Top graphs show all reads sharing nt identity >70%; bottom graphs show the subset of these reads that share nt identity >98%. The reads represent 20 populations, which were adequately sampled (abundant), combined together (Panel A; n = 411790 reads), a single representative population of these 20 abundant populations that appears to be too clonal to assess the intraspecies gap, e.g., most reads showing nt identity >99% (Panel B; n = 23109), and another single population (of the 20 abundant populations) that, while the ANI gap is not clear when all reads belonging to the population were assessed (Panel C; 16817 reads), the gap becomes evident when the analysis is limited to the rpoB-carrying reads (Panel D; 50 reads). MAG A represents a novel genus of the *Flavobacteriaceae* family and MAG B a new member of *Schleiferiaceae* affiliated at the species level with GTDB’s taxon s__UBA10364.

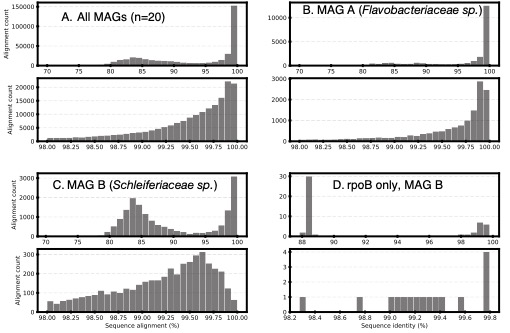

**Supplementary Figure S5. Enrichment of hypothetical and mobile genes among the genes that differ between more closely related genomes.** The graph shows the functional annotation of the gene-content differences among pairs of genomes (y-axis) plotted against the ANI value of the two genomes (x-axis). Genes were predicted using Prodigal with default settings for 16718 genomes from 279 species (2). For each species, genes were clustered with MMseqs2 at 90% amino acid sequence identity using the following parameters “--min-seq-id 0.90 --cov-mode 1 -c 0.5 --cluster-mode 2 --cluster-reassign” (3). For each species, a binary matrix was generated from MMSeqs2 output with genomes as columns and gene clusters as rows. A one indicates a gene cluster is represented in a genome. Gene differences were tabulated for genome pairs from this matrix and assigned to the corresponding ANI bin. Representatives of each gene cluster were annotated with EggNog Mapper using default settings (4). The Mobile category was generated using a keyword search of the annotation’s description column using the keywords: transposase, phage, integrase, viral, plasmid, integron, and transposon. The Metabolism category was created using the assignments from the COG_cat column: C, G, E, F, H, I, P and Q. The COG_cat value of S was reported as Conserved Hypothetical. Any gene not receiving an annotation is reported as Hypothetical. All other gene assignments were reported as Other. The number (percentage) at the top of each bar indicates the percent of total gene differences in that ANI bin; the numbers on the series indicate the fraction of the total gene content differences that the series made up.

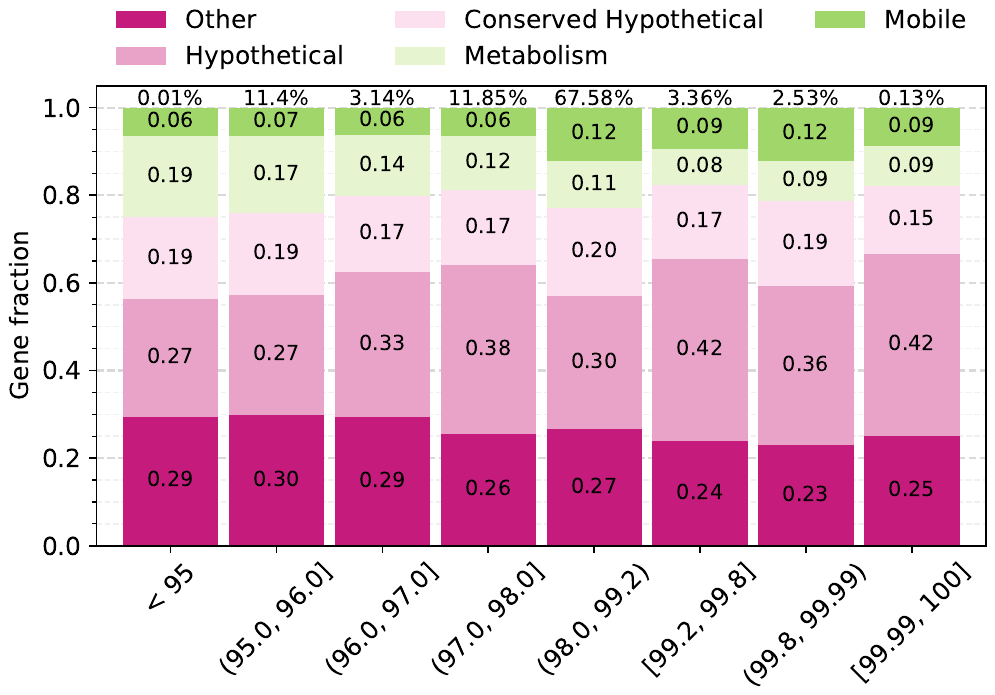

**Supplementary Figure S6. Summary statistics of the comparison of the 99.5% ANI threshold to available clonal complexes of *E. coli***. The lines represent the four most abundant STs in terms of number of genomes assigned to them (see key); these STs included ~60% of the total *E. coli* genomes evaluated and are the same as those shown in Fig. 4. F1 statistic (left panel) and Accuracy (right panel) were estimated as shown at the bottom of Table S1. Note the increase in accuracy and F1 around 99.5% ANI, consistent with the gap in ANI value distribution in the 99.2-99.8% range.

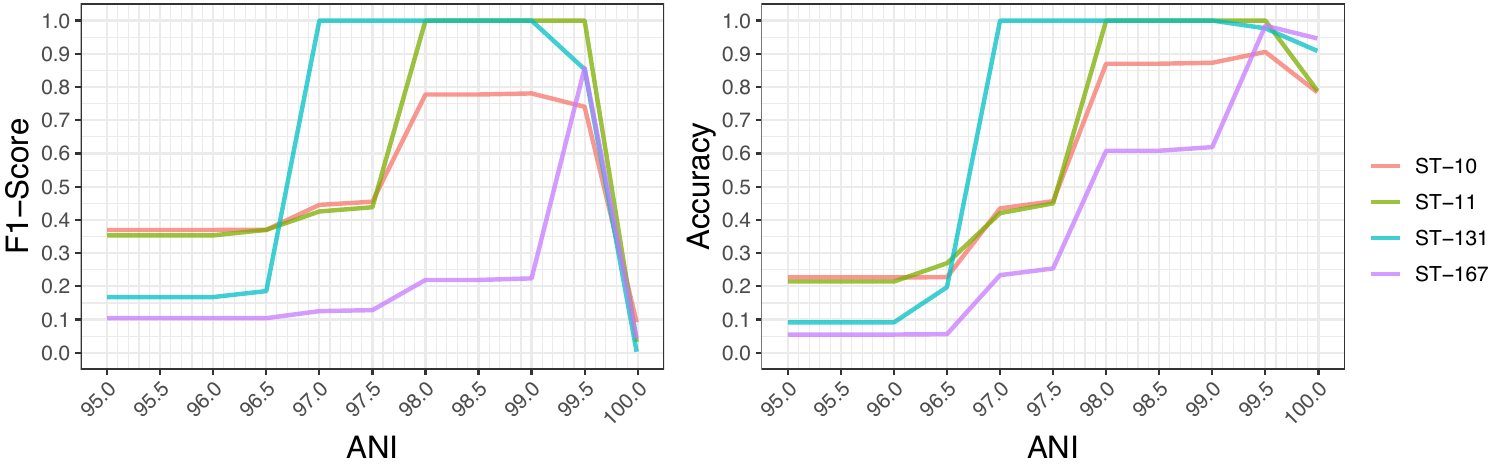
